# C12ORF57: a novel principal regulator of synaptic AMPA currents and excitatory neuronal homeostasis

**DOI:** 10.1101/2025.01.08.632037

**Authors:** Ruiji Jiang, Malek Chouchane, Kok Siong Chen, Ching Moey, Jens Bunt, Jiang Li, Fowzan Alkuraya, Eman Alobeid, Linda Richards, Erik Ullian, Elliott Sherr

## Abstract

**Objective:** Excitatory neuronal homeostasis is crucial for neuronal survival, circuit function, and plasticity. Disruptions in this form of homeostasis are believed to underpin a variety of neuronal conditions including intellectual disability, epilepsy, and autism. However, the underlying genetic and molecular mechanisms maintaining this homeostasis remain poorly understood. Biallelic recurrent loss of function mutations in *C12ORF57*, an evolutionarily conserved X amino acid novel open reading frame, underlie Temtamy syndrome (TS)—a neurodevelopmental disorder characterized by epilepsy, dysgenesis of the corpus callosum, and severe intellectual disability.

**Methods:** Through multiple lines of inquiry, we establish that C12ORF57/GRCC10 plays an unexpected central role in synaptic homeostatic downscaling in response to elevated activity, uncovering a novel mechanism for neuronal excitatory homeostasis. To probe these mechanisms, we developed a new knockout (KO) mouse model of the gene’s murine ortholog, *Grcc10* as well as cellular and *in vitro* assays.

**Results:** *Grcc10* KO mice exhibit the characteristic phenotypic features seen in human TS patients, including increased epileptiform activity. Corresponding with the enhanced seizure susceptibility, hippocampal neurons in these mice exhibited significantly increased AMPA receptor expression levels and higher amplitude of miniature excitatory postsynaptic currents (mEPSCs). We further found that GRCC10/C12ORF57 modulates the activity of calcium/calmodulin dependent kinase 4 (CAMK4) and thereby regulates the expression of CREB and ARC.

**Interpretation:** Our study suggests through this novel mechanism, deletion of Grcc10 disrupts the characteristic synaptic AMPA receptor downscaling that accompanies increased activity in glutamatergic neurons.

## Introduction

Neuronal excitatory homeostasis, the maintenance of neuronal excitability within a set range, is a crucial component of neuronal survival, growth and circuit development. It is a dynamic process balancing overall needs to maintain stability and integrity of underlying neuronal circuits while also allowing for graded changes to strengthen or weaken connections.^1^ While epilepsy is one key outcome when this process goes awry, disruption of genes involved in synaptic regulation underlie many neurologic disease such as Fragile X syndrome, autism, Rett syndrome, and neurodegenerative disease.^2,3^ To date, many individual genes have been associated with synaptic homeostasis yet, the underlying molecular mechanisms driving this homeostasis remain poorly understood. Our discovery of the central regulatory role for *C21ORF57* underscores how critical C12ORF57 is as a regulator of neuronal and synaptic homeostasis. Recent linkage and sequencing analyses have identified a recurrent autosomal recessive mutation in the start codon (c.1A>G, p.1M>V) of *C12ORF57* as the cause of Temtamy syndrome (TS), an autism and intellectual disability (ID) disorder characterized by medically intractable seizures and dysgenesis of the corpus callosum.^4–8^ This syndrome was initially reported in Middle Eastern populations but subsequently observed in other countries.^5,9–11^ This mutation is prevalent throughout the Middle East and is the single most common cause of recessive ID and developmental delay in Saudi Arabia, accounting for greater than 1% of intellectual disability with genetic cause in these cohorts.^4,5^ *C12ORF57* is a three exon gene encoding a highly conserved 126 amino acid, 13 kD protein that is abundantly expressed in human fetal and postnatal brain.^4,8^ Despite the widespread nature of this mutation among affected populations and evolutionary conservation suggesting an important biological function, currently there is little known about its role in normal neuronal development and homeostasis. This is because C12ORF57 lacks homology to any known vertebrate protein, making it difficult to predict its function (**Supplemental Figure 1**).^4^ To determine the function of this novel gene, we generated and analyzed the consequences of the homozygous knockout model of its murine ortholog, *Grcc10,* with which it shares 99.2% amino acid identity (**Supplemental Figure 1**).

The *Grcc10* knockout (KO) mice recapitulate many of the phenotypes found in human TS patients, notably epilepsy and corpus callosum anomalies, creating a valuable model to study this gene’s function. Using biochemical (including yeast-2-hybrid interactions) and electrophysiological assays, we demonstrate the mechanisms by which C12ORF57/GRCC10 regulates synaptic excitability. We analyze its interaction with the neuronal homeostasis protein, calcium calmodulin dependent kinase IV (CAMK4), as well as its regulation of CREB, Fos, and ARC, which are implicated in the synaptic scaling of AMPA receptors.^1^ Collectively, our data show for the first time, that C12ORF57 is a central regulator of synaptic homeostasis, revealing a novel molecular mechanism of neuronal excitatory tuning.

## Materials and Methods

### Animal maintenance and generation of G*rcc10* mouse lines

All procedures of animal maintenance and experiments were in accordance with the regulations of and approved by IRB at the University of California San Francisco and in compliance with relevant federal and local laws. All animal breeding and procedures performed in the Richards lab were approved by The University of Queensland Animal Ethics committee and performed in accordance with the Australian Code of Practice for the Care and Use of Animals for Scientific Purposes. Generation of the mouse *Grcc10<tm1.1(KOMP)Vlcg>* line on a C57BL/6J background by the Sherr lab was previously described.^12,13^

Embryos and postnatal pups were bred from wildtype CD1 time-mated dams at The University of Queensland. The detection of vaginal plug in the following morning after mating was designated as embryonic day zero (E0). Pups are at postnatal day (P0) on day of birth. Tissue collection and fixing was as previously described (9), then post-fixed with 4% paraformaldehyde (ProSciTech) for 2 to 4 days and stored in 1x Dulbecco’s phosphate buffered saline (PBS, Lonza) with 0.2% sodium azide (Sigma Aldrich).

### Primary Fibroblast Derivation and RT-PCR

Primary fibroblasts were derived from 10mm skin punch biopsies of C12ORF57 patient as previously described by Kisiel et al.^14^ All fibroblasts were below passage number 3. RNA was extracted using PureLink RNA mini system (ThermoFischer 12183018A). cDNA was generated from RNA samples using superscript IV one Step RT-PCR system (ThermoFischer, 12594025). RT PCR was run using the following primers: C12ORF57 F:5’-GGATAACGCCTGCAACGACATG-3’ R: 5’-CTTCGTAGGACTTGACCAAGCG-3’; GAPDH: F: 5′-GTCTCCTCTGACTTCAACAGCG-3′; R: 5′-ACCACCCTGTTGCTGTAGCCAA-3′

### Kainic Acid Treatment

Postnatal day 21 genotyped mice received 20 mg/kg intraperitoneal kainic acid (Sigma-Aldrich K0250) solution dissolved in saline or saline vehicle solution using sterile technique. An observer blinded to mouse genotype recorded time to each Racine stage.

### C12ORF57 Antibody generation

C12ORF57 peptide was generated using a full-length cDNA of human C12ORF57 conjugated to N-terminal GST (NCBI9606) produced in BL21(DE3) competent *E.coli.* and purified against a glutathione agarose (GE Lifescience 17075601). Rabbit polyclonal antibody was produced using ThermoFisher Custom Polyclonal Antibody production.

Antibody showed specific bands on western blot at 1:200 dilution in both CMV-C12ORF57-FLAG at expected 17 kD weight and in Grcc10 WT mouse brains and not in Grcc10 KO brains (**Supplemental Figure 2**).

### Cellular Fractionation

Whole mouse brains were Dounce homogenized in fractionation buffer (20 mM HEPES, 10 mM KCl, 2mM MgCL2, 1 mM EDTA, 1 mM EGTA, 1mM DTT) followed by passage through a 27-gauge needle. The nuclear fraction was separated by centrifugation at 720 xg for 5 min. Cytoplasmic fraction was separated at 10,000 xg for 5 min.

### Western Blot

Tissue was homogenized with a Dounce homogenizer in RIPA buffer for whole cell lysates, and lysates were run on a 8-20% gradient SDS page. The blots were blocked with Odyssey Li-Cor blocking buffer (Licor). Primary antibodies were followed by an IR fluorescent-conjugated goat anti-mouse and anti-rabbit IgG (1:10 0000, Licor). Quantification and IF intensity analysis was done using Odyssey LiCor Image Studio Software v.5.2.5.

### Antibodies

The following primary antibodies were used (Species, Company, Clone, dilution): CAMK4: Invitrogen 8C5B8 (Mouse, MA5-17038,1:1000), pCAMK4 Thr200 (Rabbit, Invitrogen, PA5-37504, 1:1000), pCREB: (Mouse, CST 87G3 #9197, 1:1000), CREB (Rabbit, CST, 48H2, 1:1000), GFP (Chicken, A10626, Invitrogen,1:1000), GAPDH (CSF, Rabbit, 1:1000), Lamin AC (CST, #2032, 1:1000) Arc (Proteintech, Rabbit, 16290-1-AP, 1:1000), AMPA (6C4, Invitrogen, 32-0300), and CTCF (CST, 28995, Rabbit,), Caveolin-1 (Rabbit, CST, D46G3, 1:1000) The following secondary antibodies were used: IRDye680RD Goat anti-Rabbit IgG (H+L) 1:10,000 (Li-cor, 926-68071), IRDye680 Donkey Anti-Chicken (Li-cor, 926-68075), and IRDye800 Goat anti-mouse IgG (Li-Cor, 926-32210).

### Co-immunoprecipitation

Mouse brains were homogenized in NP40 lysis buffer (1% NP40, 240 mM NaCl, 5 mM EDTA 50 mM Tris ph7.4). 1µg of rabbit anti-C12ORF57 antibody was bound to 2µl of agarose resin and incubated in 1.5 µg of total lysate overnight and co-immunoprecipitated using Pierce Co-Immunoprecipitation kit (ThermoFisher, 26159). Protein was dissociated using low pH Elution buffer (ThermoFisher, 21104).

### Comparative Pull-down experiments

HEK293 cells were co-transfected with C12ORF57-DYKDDDK and CAMK4 constructs in a pCMV backbone. Cells were lysed with 1% NP40 lysis buffer in 1x PBS.

1 ug of rabbit Anti-DYKDDDK primary was bound to 20 µl of agarose resin (Pierce, 26159) and incubated in 1.5 ug of total protein overnight. Western blotting was performed as described above and bands were quantified using an Odyssey Licor system as described above.

### Hematoxylin and eosin staining

Hematoxylin and eosin (H&E) staining using standard H&E protocols was performed on P0 Grcc10 knock-out mouse brains. Dissected brain tissues were sectioned on vibratome (Leica) at 50µm thickness in sagittal and coronal orientation.

### In situ hybridization

In situ hybridization was performed as described in Moldrich et al, 2010.^15^ Two sets of Grcc10 riboprobes were generated in-house using the primers corresponding to those used in the Allen Mouse Brain Atlas (2015 Allen Institute for Brain Science http://mouse.brain-map.org/experiment/show/386251, Forward primer 1: 5’ ACTCAAGCACCGAGTGGC 3’, Reverse primer 1: 5’ CCTCTGGAACACAAGGGC 3’) (Forward primer2: 5’ GCTTCCTTTCTGCCTCCTCT 3’, Reverse primer 2: 5’ CACAGCAGCAGCACACATAC 3’). and primers designed with Primer3Plus software. These primers were used to amplify a 459 or 519 base pair fragment, respectively, from mouse cortex cDNA. These fragments were purified and cloned into pGEM-T Vector System (Promega). The plasmids were then linearized (NotI or SacII restriction enzyme, New England BioLabs), purified (PCR Clean up Kit, Qiagen), transcribed (Sp6 or T7 RNA Polymerase, New England BioLabs) and digoxigenin-labelled (DIG RNA Labelling Mix, Roche) or fluorescein-labelled (Fluorescein RNA Labelling Mix, Roche) to generate the riboprobes.

### *In Vitro* CAMK4 assay

CAMK4 kinase activity was measured using ADP Glo System (Promega, V6930). CAMK4, calmodulin, and reaction buffer were supplied by CAMK4 *in vitro* Kinase system (Promega, V2951). C12ORF57-FLAG was transfected into HEK293 cells and purified from whole cell lysate through immunoprecipitation on Anti-DYKDDDK Affinity resin (Genescript, L00432). CAMK4, C12ORF57, and purified recombinant PP1CA (Origene, TA808819) were mixed in equimolar amounts with 25 µM ATP and reaction luminescence was measured on a Tecan Spark with an integration time of 1 sec.

#### Primary Hippocampal Neuronal Culture and Transfection

Embryos were removed from pregnant dams at embryonic day 18 (E18). Quick genotyping of the mice was performed by PCR analysis of DNA, extracted from the tail biopsies. Tails were digested in KAPA Express Extract Kit and then template DNA was added to PCR mix containing: 1X KAPA2G Robust HotStart ReadyMix, and *Grcc10* primers (Forward*: 3’-* GCGGACCTCTCTGGGATG-5’ Reverse: 3’-GCAGCACACATACGAGAACC-5’)

After genotyping, hippocampi were dissected in ice-cold Hanks’ balanced salt solution (HBSS; Gibco) and were incubated in trypsin/EDTA (Gibco BRL) at 37 °C for 25 min and subsequently transferred to trypsin inhibitor (Worthington, LS003570) at room temperature for 5 min. After trituration in DMEM/10% FBS solution, cells were counted and plated on poly-L-lysine (Sigma, P4707) and Laminin (Gibco, 23017015) pre-coated cover glass (Harvard Apparatus) at a density of 40.000 cells per 10 mm cloning rings. Cells were maintained in Neurobasal medium (Gibco) containing 1% heat-inactivated FBS, 1x B27-supplement (Gibico), 1X Glutamax (Gibco), 15 mM NaCl. The medium was partially exchanged for fresh every 7 days. On days 3-5 (days in vitro, DIV), mitotic inhibitor 5-fluoro-2′-deoxyuridine (FUDR, 10 μM) was added to control glial growth.

For the transfected experiments, on day 7, *Grcc10+/+* neurons were infected by custom lentiviral particles produced from CW307508 (Origene) with C12ORF57-P2A-GFP under control of a synapsin 2 promoter (Origene, Catalog No: CW307508V) at a concentration of 1:100 GFP under *Syn2* (Origene, Catalog No: RC218378LF) or CAMK4-P2A-GFP (Origene Catalog No: MR207505L4V).

#### In vitro patch clamp recordings

Patch clamp measurements were conducted at DIV 14-17. Before recordings, Neurobasal medium was replaced by HEPES ACSF, containing (in mM): NaCl, 140; KCl, 2.5; HEPES, 10; MgCl2, 2; CaCl_2_, 2; and dextrose, 10. pH was adjusted to 7.3 with NaOH. mOsm/kg was set to 300. Cultures were visualized under upright microscopy (Olympus, BX51WI) and cells with pyramidal morphology were identified by ×40 water-immersion objective (Olympus) and IR-2000 (DAGE-MTI) camera GFP was identified with 488 nm LED pulse. Whole-cell patch clamp recordings were done from the soma of the pyramidal hippocampal neurons using Axopatch 200B amplifier (Molecular Devices, USA). Cell capacitance and access resistance were monitored throughout the experiments. Signals were acquired with a DigiData-1440A digitizer controlled by pCLAMP-11 software (Molecular Devices, USA). Signals were sampled at 10 kHz and low-pass filtered at 2 kHz. Borosilicate glass pipettes (4–5 MΩ) were pulled with a horizontal puller (P-1000, Sutter Instrument).

Miniature excitatory currents (mEPSCs) were isolated from the GABAergic transmission by applying 50 µM GABA_A_ blocker picrotoxin in ACSF. To capture mEPSCs, voltage-gated Na+ channels and NMDA receptor-mediated currents were blocked with 1 µM TTX and 50 µM D-AP5, respectively. Patch pipettes were filled with Cs-based intracellular solution (in mM): CsMeSO3, 135; CsCl, 6; HEPES, 10; phosphocreatine, 10; EGTA, 0.6; Mg-ATP, 3; Na-GTP, 0.5; QX314-Br, 2. pH was set to 7.35 by CsOH and osmolarity to 305 - 310 mOsm/kg. mEPSCs were measured from the holding membrane potential of -70 mV. NMDA receptor-mediated currents were recorded using the same external solution, without magnesium and D-AP5, and with the addition of NBQX to block AMPA currents (10 µM, Tocris). GABA receptor-mediated currents were recorded using external solution without picrotoxin, but with D-AP5, TTX and NBQX in its place. Data analyses were done using Easy Electrophysiology software (Easy Electrophysiology Inc) utilizing a template modeling process for mEPSC detection.

#### C12ORF57 HEK293 Knockout

sgRNA were generated with the sequence 3’-GATAACGCCTGCAACGACAT-5’ and inserted into pCAS-Guide-EF1a-GFP vectors (Origene). HEK293-17T cells were transfected with Lipofectamine 3000 (ThermoFisher, L3000001) and the transfection efficiency was determined via immunofluorescence. Lines were picked and maintained with DMEM+10% FBS containing puromycin at 1 ug/ml.

#### Golgi Cox Staining and dendritic spine density

P18 mouse brains were fixed in formalin and frozen sectioned. Golgi-Cox staining was performed on these neurons utilizing the FD rapid GoligStain Kit (FD Neurotechnologies Inc PK401A). Neurons were chosen at random from slices of mouse cortex and imaged on a Nikon T1 inverted microscope. Dendritic spines were counted and analyzed using ImageJ software plug in Dendritic spine counter.

#### Neuronal Surface Staining

Live cultured KO and WT neurons were incubated with a primary antibody directed against the extracellular domain of GluA2. Incubation was conducted at room temperature to slow down endocytosis of the antibody as shown previously.^16,17^ After incubation with the primary antibody, cultures were washed and fixed for standard immunostaining. GluA2 positive puncta were imaged using confocal microscopy. Their number and fluorescence intensity were quantified over 50 µm of neuronal processes using ImageJ software.

#### Bicuculline Inhibition

Surface AMPA receptor expression after induction of a chronic increase in postsynaptic activity was assessed by measuring GluR2 surface expression before and after treatment of neurons with bicuculline for 48 hours, as described previously.^12,17^

#### Statistical analysis

Comparisons between Genotypes:

Analysis of Variance (ANOVA): T-test or Mann Whitney or ANOVA were appropriate was conducted to assess the overall differences between genotypes. When possible, all measurements were done by blinded observers. All t-test were two-tailed.

In case of significant differences identified by ANOVA, post hoc tests (Tukey’s Honestly Significant Difference test, Sidak’s multiple comparison test) were performed to determine specific pairwise differences between groups.

Data represent +standard deviation.

Statistical software

All statistical analyses were performed using Prism version 10.0.0. The significance level (α) was set at 0.05 for statistical tests. Statistical differences are indicated in the figures and tables using the following symbols: \**P* ≤ 0.05, \*\**P* ≤ 0.01, \*\*\**P* ≤ 0.001, \*\*\*\**P* < 0.0001.

## Data Availability

The data that support the findings of this study are available from the corresponding author, upon reasonable request.

## Results

### Grcc10 loss of function in mice recapitulates Temtamy syndrome phenotypes

When examined by RT-PCR, the *C12ORF57* transcript was not detected in primary fibroblasts from patients homozygous for the most common c.1 A>G variant, suggesting that their phenotype was consistent with a complete loss of function (LoF) (**Figure 1A**). ^4,5^ To model this homozygous inactivation of *C12ORF57* in human disease, we generated a transgenic mouse line (*Grcc10<tm1.1(KOMP)Vlcg>*) carrying a beta galactosidase-containing cassette that constitutively disrupts the *Grcc10* gene (*Grcc10* KO). *Grcc10* +/+ (WT), +/- (Het) and -/- (KO) littermates were born with the expected 1:2:1 Mendelian ratios from heterozygote pairs (X=0.014, dF=2, p=0.99, Chi-Square, n=154) **(Figure 1B**). *Grcc10 -/-* pups had high postweaning mortality compared to WT and Het littermates as early as postnatal day 3 (P3) (X=.014, df=2, p=0.00013, Chi-Square, n=91), with the majority of animals dying before P2 (**Figure 1B**). *Grcc10* KO mice that survived until weaning age showed significantly diminished physical growth compared to their WT and Het littermates (11.71±0.11g vs. 4.15±0.99g, respectively, p<0.00001 t=33.07, df=8.18, N=5, Welch’s T-Test), suggesting that *Grcc10* is also crucial for health and survival (**Figure 1C**).

**Figure 1.**
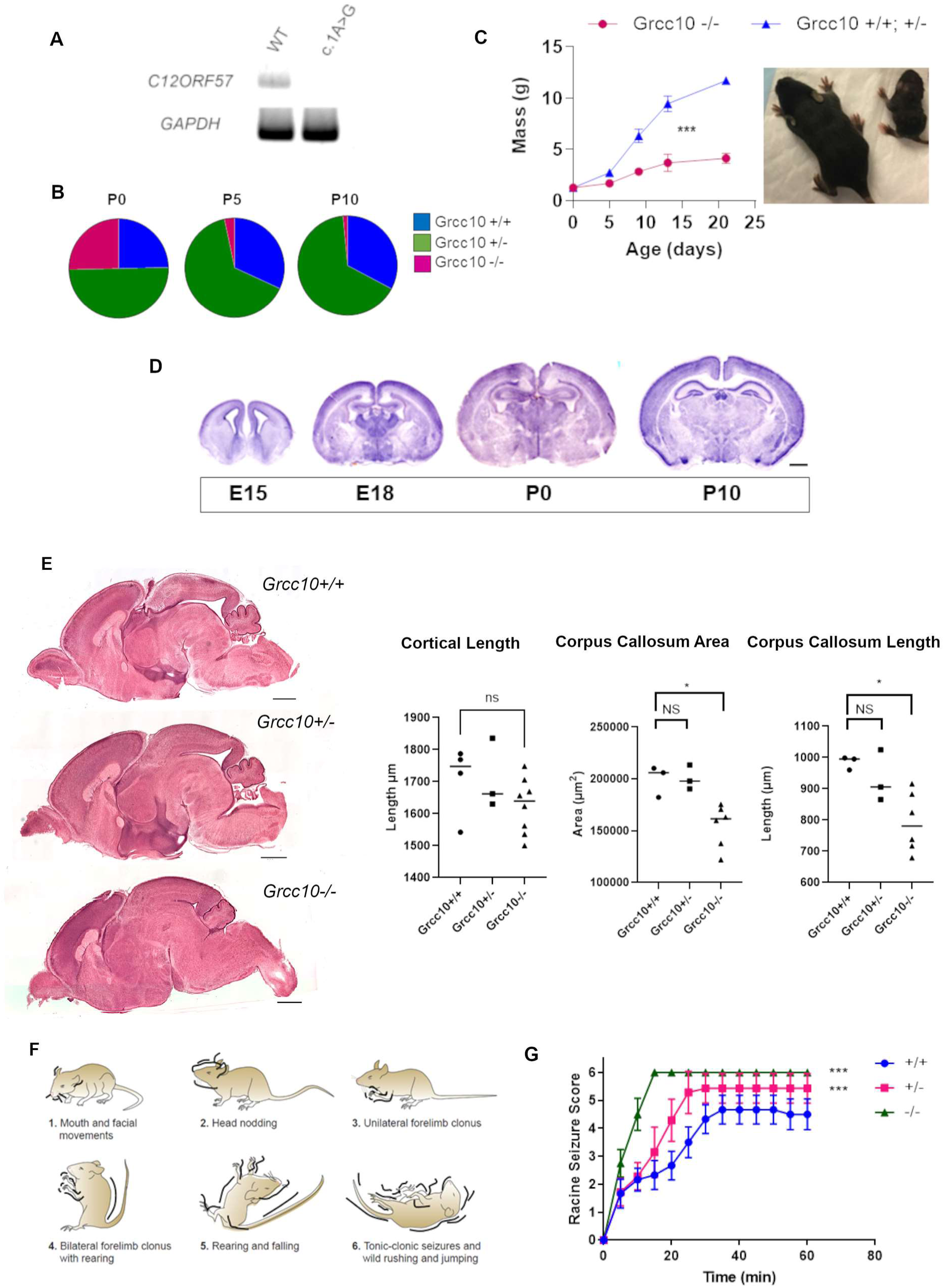
Grcc10-/- Mice Recapitulate the Temtamy Syndrome Phenotype. **A)** RT PCR of C12ORF57 from fibroblasts derived from a human patient with homozygous c.1 A>G mutations in C12ORF57 see loss of transcript when compared to wild type (WT) controls. **B)** Ratios of surviving *Grcc10* genotype mice are shown at ages P0, P3, and P21 (N=154, p=0.99, Chi-Square). **C)** Mean body mass (g) for ages P0, P5, P9, P13, and P21 (N=5 for KO mice N=14 for WT and heterozygotes, p>0.0001 ANOVA) **D)** *In situ* hybridization of C12ORF57 shows diffuse widespread expression of *Grcc10/C12ORF57* throughout all brain regions throughout embryonic development. **E)** Representative images of midsagittal slices from *Grcc10+/+, Grcc10+/-* and *Grcc10+/+* mouse brains (Scale Bar 1000µm) along with graphs of cortical length (p=0.1127, ANOVA with Tukey post-hoc test), corpus callosum length (p=0.017, ANOVA with Tukey post-hoc test) and corpus callosum cross sectional area cross-sectional area (p=0.0058, ANOVA with Tukey’s post hock). N=6 for KO, N=3 for WT, N=3 for Het. Median represented by horizontal line **F)** Pictographic of Racine stages **G)** Average Racine stage (x-axis) of mice in time post kainic acid injection (y-axis) for *Grcc10+/+* (blue, N=6), *Grcc10+/-* (pink, N=7), and *Grcc10-/-* (green, N=4) (p=0.00162, ANOVA with post hoc Tukey test). Statistical significances are indicated by the following symbols \**P* ≤ 0.05, \*\**P* ≤ 0.01, \*\*\**P* ≤ 0.001, \*\*\*\**P* < 0.0001.

Since our previous work associating *C12ORF57* with ID, ASD, and epilepsy, and its high expression in human fetal brain tissue strongly suggests a crucial role in normal brain development, we first investigated *Grcc10* expression in early mouse brain development using *in situ* hybridization on late embryonic days (E)15 and 18 (E15, E18) and P0 and P10.^5,8^ *Grcc10* is broadly expressed in many brain regions at all examined ages. At E15, it is specifically localized to the cortical plate, glial wedge, and dorsal ventricular zone. At E18, we observed higher expression in the cortex, hippocampus, and thalamus that was maintained until P10 (**Figure 1D**). Interrogation of Allen Brain Atlas cell and region-specific RNA-seq also shows global expression across multiple brain areas and neuronal cell types (**Supplemental Figure 3**). The widespread expression of *Grcc10* in the developing cortex suggests that *Grcc10* may play an early and central role in cortical development. In addition, as previous studies showed that disrupted cortical lamination is linked to ASD, ID and epilepsy we examined whether this was the case in *Grcc10* KO mice by performing immunohistochemistry on P0 murine cortical slices using previously validated upper and lower-layer markers *Satb2* and *Ctip2* respectively.^18^ We observed no difference in the number of Satb2 WT (1403.00±89.09 cells/HPF) versus KO brains (1498.20±40.14, p=0.36, t=0.95, df=10, Welch’s tailed T-test) or Ctip2 neurons in WT (610.43±31.00) versus KO brains (633.17±41.03, p=0.76, t=0.32, df=11, Welch’s T-test) or the pattern of lamination of WT and *Grcc10* KO cortices, suggesting that *Grcc10* loss of function does not impair gross cortical architecture or layering (**Supplemental Figure 4**).

Our *Grcc10* -/- mice share a shortened corpus callosum structure similar to what is found in patients with Temtamy syndrome. In P0 KO mice, corpus callosum length was shorter (792.81±39.15 µm, p=0.017, F=6.41, df=2. N=6, ANOVA) when compared to their WT (984.53±12.23µm, p=0.023, df=9, Tukey post-hoc test), and heterozygous littermates (931.89±47.73µm) though this was not statistically significant (p=0.092, q=3.391, DF=9, Tukey post-hoc test). The corpus callosum cross sectional area in these P0 KO mice was also reduced (154859.22±8441.27µm^2^, p=0.0058, ANOVA, F=9.65, df=2 N=6) compared to WT (199602.27±8702.73µm^2^, p=0.012,q=3.546, df=9, N=3, Tukey post-hoc test) and Het mice (200644.93±6764.12µm^2^, p=0.010 q=3.628, DF=2, N=3, Tukey post-hoc test). Rostral to caudal cortical length between KO (1625.01±30.49µm) and WT (1706.25±56.16 µm,) or Het (1709.39±63.84 µm p=0.29, ANOVA, F=1.337, df=2, N=6) was not significantly different (**Figure 1E**).

Patients harboring *C12ORF*57 homozygous or biallelic mutant variants have a high rate of epilepsy that is refractory in many cases.^5,6^ We have observed *Grcc10* KO mice displaying spontaneous seizure activity (**Supplemental Video 1**), but their early mortality and small size prevented us from conducting EEG monitoring. To assess seizure susceptibility, we tested whether these mice are more susceptible to pharmacologically induced seizures. We performed intra-peritoneal kainic acid injection in P21 *Grcc10* KO, Het and WT mice, and scored their behaviors indicative of seizures based on the Racine scale of 1-5 (**Figure 1F**).^19–22^ All animals exhibited seizures following kainic acid administration, whereas no seizures were observed after vehicle injections. *Grcc10* KO mice reached clinically severe stages (=>5) significantly faster (9.44±1.33 min,p<0.0001, 2-way ANVOA F=46.34, F=764.66 N=4) than *Grcc10* WT (36.33±2.84 min, p<0.0001, N=6, Tukey post-hoc test) or Het littermates (20.74±1.82 min, N=5 p<0.0001, Tukey post-hoc test) (**Figure 1G**). These data suggest that *Grcc10* KO mice have higher underlying susceptibility to induced seizures, analogous to the increased rate of epilepsy seen in human Temtamy syndrome patients.

### Grcc10 loss of function causes an increase in AMPA mediated mEPSCs

We sought to determine what synaptic changes may mediate the severe seizure activity seen in our mouse model. We hypothesized that we would be able to see increased neuronal excitability through AMPA (α-amino-3-hydroxy-5-methyl-4-isoxazolepropionic acid) mediated currents based on substantial evidence from many animal models that shows that these receptors are key mediators of seizure spread and epileptogenesis.^23,24^ We measured electrophysiological parameters in *Grcc10* KO neurons by first assessing spontaneous AMPA mediated miniature excitatory postsynaptic currents (mEPSC) using whole cell recordings in isolated excitatory hippocampal neurons from E18 *Grcc10* WT and KO mice on day *in vitro* (DIV) 14-18. We observed that AMPA-mediated mEPSCs recorded from *Grcc10* KO neurons have a significantly higher mean amplitude than mEPSCs recorded from *Grcc10* WT neurons (35.81±0.90 pA vs 15.49±0.29 pA, respectively, p<0.0001, F=367, df=2, N=5 neurons per genotype, ANOVA with Tukey post-hoc test) (**Figure 2A-C**). These data support the hypothesis that *Grcc10* loss of function causes an increase in AMPA-mediated mEPSC amplitudes. In contrast, NMDA amplitude and frequency were not significantly different between WT and KO neurons (13.84±3.93 pA vs. 12.92±2.92 pA, p=0.20, t=1.27, df=301.4, N=5, Welch’s T-test) (**Figure 2D**). GABA current amplitude was also not significantly different between WT and KO neurons (13.78±3.32pA vs. 15.56±8.37 pA, (p=0.16, t=1.41, df=137, N=5 Welch’s T-test) (**Figure 2E**). To confirm that the loss of C12ORF57/GRCC10 was a necessary and sufficient cause of the increase in AMPA mEPSC activity in these cultured neurons, we sought to rescue the *Grcc10* KO neurons using lentiviral vector containing *C12ORF57-P2A-GFP* construct under *Syn2* translational control with a *GFP* only lentivirus serving as a control (**Supplemental Figure 5**). We transfected E18 *Grcc10* KO neurons on DIV 7 with both the construct and a *GFP* only vector and compared their electrophysiology profiles on DIV14-18 with WT lentiviral infected *GFP* controls. *Grcc10* KO neurons transfected with *C12ORF57* had decreased mEPSC amplitude compared to *GFP*-transfected KO neurons (18.27±0.47 pA vs. 38.93±0.56 pA, respectively, p<0.0001) and had no statistically significant difference in mEPSC amplitude compared to *GFP* transfected WT neurons (15.50±0.32 pA; p=0.57, t=0.17, df=137.2 N=4, ANOVA with Tukey post-hoc test) (**Figure 2F**). Frequency of AMPA mEPSC also decreased in *C12ORF57*-transfected compared to *GFP*-transfected KO neurons (5.46± 0.61 events/sec vs. 22.46 ±0.47 events/sec, respectively, p<0.0001, ANOVA with Tukey post-hoc test, F=85.3, df=2, N=5). *C12ORF57*-transfection rescued the phenotype in KO neurons (19.06±0.81 events/sec, p<0.0001, ANOVA Tukey post-hoc test, N=5), which were now comparable to WT neurons (5.21±1.31 events/sec p>0.999, ANOVA with Tukey post-hoc, df=2, N=5) (**Figure 2G**). These data suggest that C12ORF57 specifically modulates post-synaptic AMPA current and is sufficient to rescue the excessively increased excitability phenotype seen in *Grcc10* KO neurons and is necessary for normal neuronal electrophysiology.

**Figure 2.**
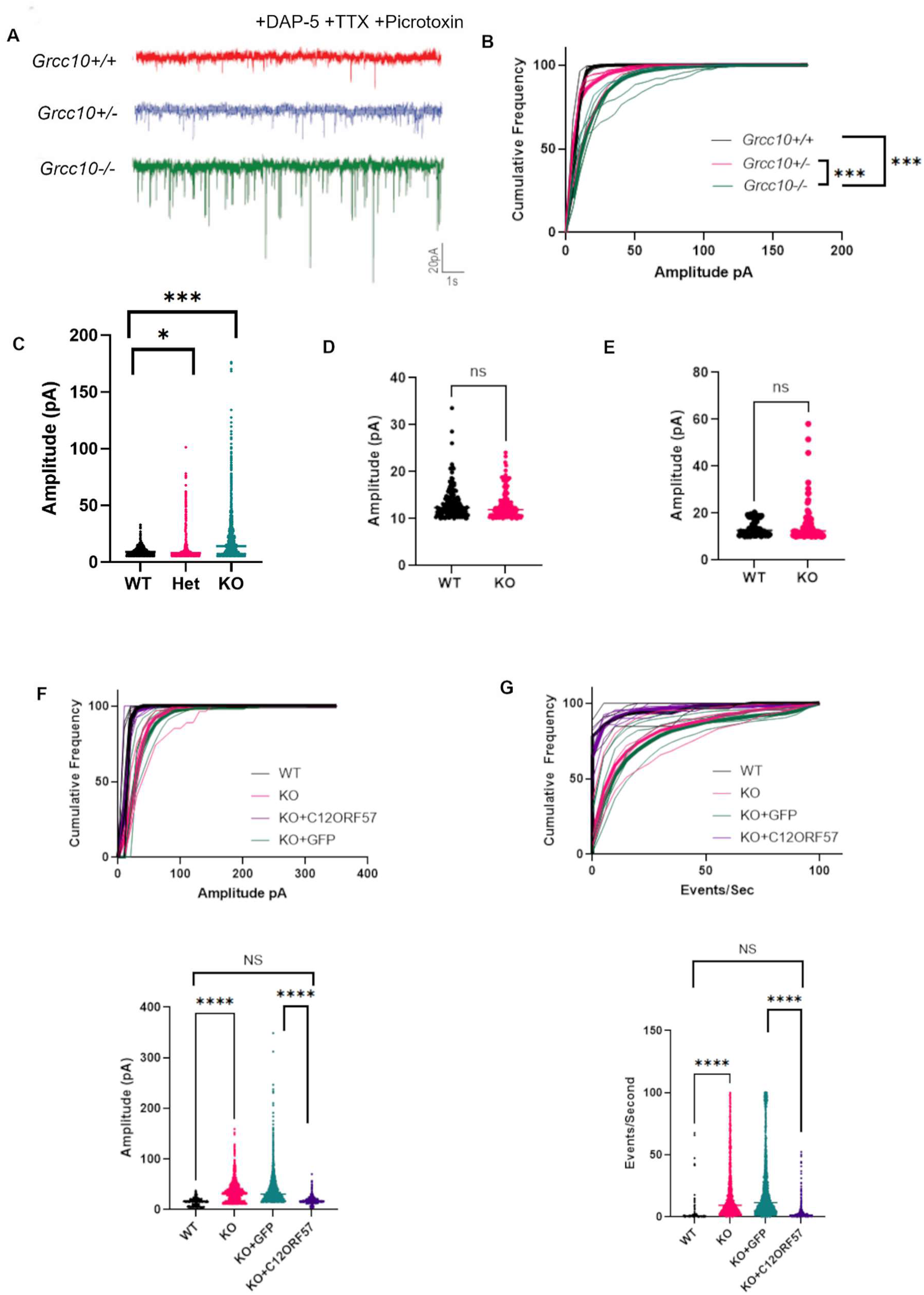
Grcc10 -/- neurons have increased mEPSCs amplitude and frequency. **A)** Representative AMPA mEPSC tracings of *Grcc10* +/+ (WT) neurons (top), *Grcc10* +/- (Het) neurons (middle) and *Grcc10* -/- neurons (bottom). **B)** Graph of cumulative frequency (x-axis) of mEPSC against amplitude in DIV 17 pyramidal neurons. The mean curve of each group is bolded. **C)** AMPA mEPSC amplitudes (pA) (p<0.0001, N=5 for each group, ANOVA) **D)** NMDA mEPSC amplitudes (pA) with median represented by horizontal line. (p=0.20 N=5 Welch’s T-test) **E**) GABA mEPSC amplitude with median represented by horizontal line (p=0.16, N=5 for each group, Welch’s T-test) **F and G)** Upper panels: Cumulative frequency of events plotted against event amplitude (left panel) and the time between events (events/second) (right panel) in DIV17 pyramidal neurons including C12ORF57 transfected rescue (purple) and GFP Control (green). The combined mean of group is represented by corresponding bolded line. Lower panels: Mean amplitude of AMPA mEPSC (left panel) and mean events/second of AMPA mEPSC (right panel). For (**F**) and (**G**), N=5 for each group, ANOVA with Tukey post-hoc test). Statistical significances are indicated by the following symbols \**P* ≤0.05, \*\**P* ≤ 0.01, \*\*\**P* ≤ 0.001, \*\*\*\**P* < 0.0001.

### Loss of *Grcc10/C12ORF57* leads to an increase in surface AMPA receptor localization and impairs excitatory homeostatic synaptic down-scaling after blocking GABA receptors

Given our findings of increased AMPA current in *Grcc10-/-* neurons we hypothesized that loss of *Grcc10/C12ORF57* would result in increased GluA2 surface expression. AMPA receptors are post-synaptic heterotetrameric assemblies of two homologous subunits (GluA1-4), with compositions containing two GluA2 subunits being the most predominant in hippocampal pyramidal neurons and mediating the majority of fast excitatory synaptic transmission in the brain. We stained for membrane bound GluA2 subunits on isolated DIV 17 *Grcc10* WT an KO) pyramidal hippocampal neurons and quantified the fluorescence intensity of GluA2 positive puncta per 50 µm of neuronal processes. We observed an increase in the number of GluA2 puncta (83.72±23.10%,(p=0.0024, t=3.62, df=15.32, N=10 per genotype Welch’s T-test) and an increase in puncta intensity (102.43±28.66%, p=0.0008, t=3.572 df=51.84, N=19 WT, N=35 KO, Welch’s T-Test) in *Grcc10* KO neurons compared to WT neurons (**Figure 3A**).

**Figure 3.**
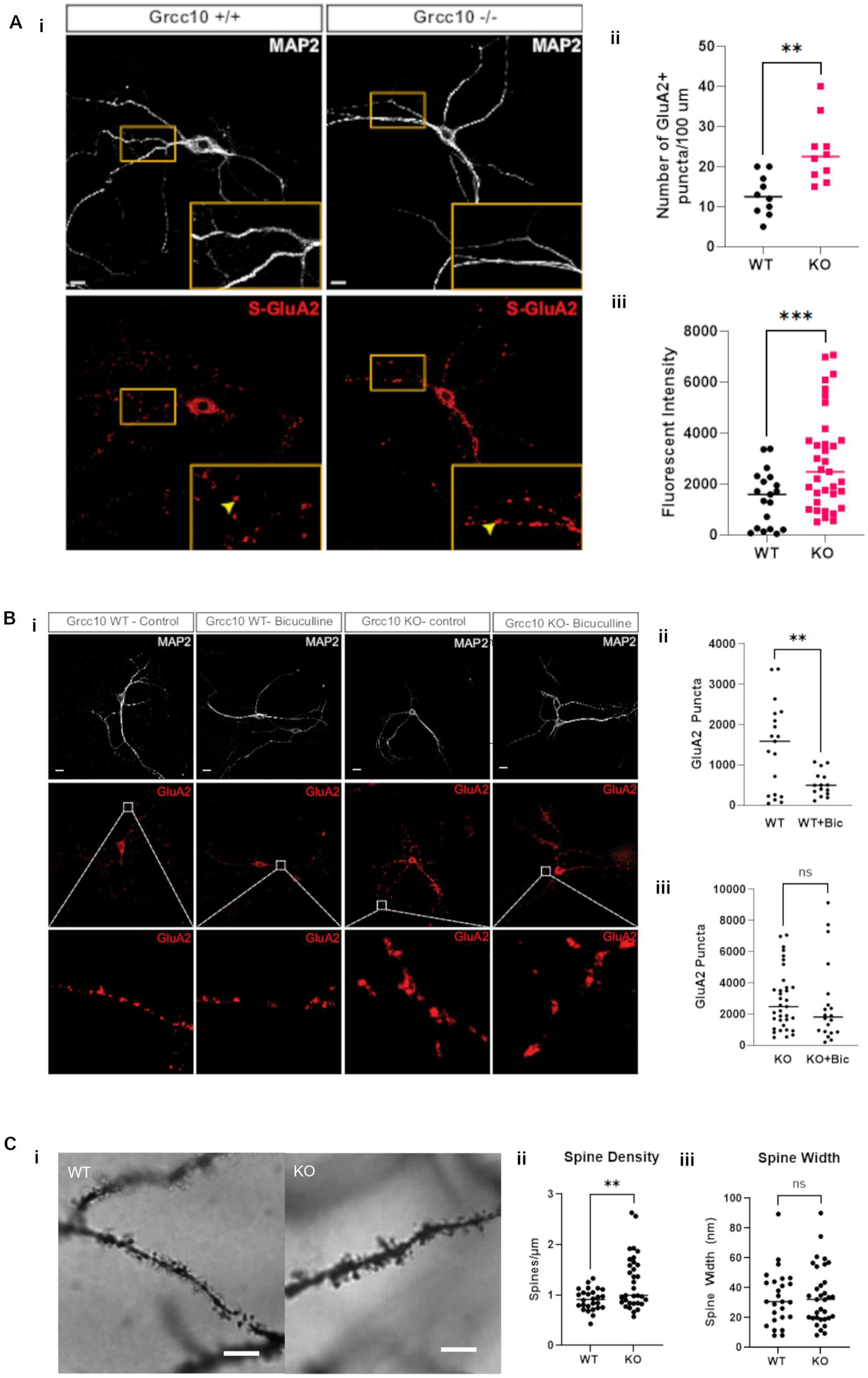
Loss of Grcc10/C12ORF57 disrupts normal GluA2 expression in primary murine neurons. **A)** (i) Representative images of surface staining of MAP (white, upper panels) and GluA2 (red, lower panels) in *Grcc10* +/+ and *Grcc10-/-* neurons. Insets 2x magnification. Scale bar 20 µm. Mean GluA2 fluorescent puncta/100µm (ii) (p=0.0024, N=10 both genotypes Welch’s T-test), and mean puncta intensity (ii) (p=0.0008, N=19 WT, N=35 KO, Welch’s T-Test) Error bars represent SEM. **B)** (i) Sample images of DIV 17 hippocampal primary neurons stained for GluA2 (red) and MAP2 (white) on both *Grcc10* WT and *Grcc10* KO neurons with and without bicuculline treatment. Scale bar 20 µm. Insets 10x magnification (ii) GluA2 puncta/100 µm with and without bicuculine, with median represented as horizontal line (p=0.0041, N=19 WT Ctl, N=15 WT+bicuculline treated Welch’s T-test. Plot of GluA2 puncta/100µm on KO neurons (p=0.68, T=0.41, df=53, N=20 KO Ctl, N=35 KO+bicuculline treated. Welch’s T-Test). Scale bar 20 µm. **C)** (i) Representative Golgi-Cox-stained dendrites from *Grcc10* WT and KO cortical neurons (scale bar 5µm). (ii) Plot of spine density (p=0.0023, N=33 per group, Welch’s T-Test) and (iii) spine width (p=0.83, N=33 per group, Welch’s T-Test). Medians represented by horizontal line in all graphs. Statistical significances are indicated by the following symbols \**P* ≤ 0.05, \*\**P* ≤0 .01, \*\*\**P* ≤ 0.001, \*\*\*\**P* < 0.0001.

We also examined whether Grcc10 KO neurons exhibit impaired downregulation of AMPA currents following increased excitation, as predicted by the decreased CAMK4 activity observed in KO mice. To do this, we measured surface AMPA receptor expression after inducing a chronic increase in postsynaptic activity by treating neurons with bicuculline (a GABA receptor antagonist) for 48 hours and then measuring GluA2 surface expression.^25^ We observed that, as expected, bicuculline treatment caused a significant decrease in surface GluA2 expression in WT neurons (-907.00±293.20 puncta/HPF, p=0.0041, t=3.09, df=32, untreated N=19, treated N=15, Welch’s T-test). In contrast, bicuculline treatment had no significant effect on GluA2 in *Grcc10* KO neurons (-255.0±616.9 puncta/HPF, p=0.68, T=0.41, df=53, N=19 untreated, N=15 bicuculline treated, Welch’s T-Test) (**Figure 3B**). We also observed an increase in dendritic spine density in *in situ Grcc10* KO cortical neurons (1.26+/-0.53 spines/µm) compared to WT cortical neurons 0.91+/-0.22 spines/µm, p=0.0023, t=3.185, df=57, N=33, Welch’s T-Test) (**Figure 3C**), though spine width was not statistically different between the two neuron types (33.53+/-18.57 vs 34.53+/-19.75, p=0.83, t=.198, df=57, N=33, Welch’s T-test). These findings suggest that GRCC10 loss of function increases dendritic spine density and GluA2 surface expression and impairs homeostatic synaptic down-scaling of GluA2 and results in higher levels of AMPA expression at synapses both at baseline and under conditions of acute increases in neuronal activity. This data further supports our observation of increased AMPA current in *Grcc10-/-* mouse neurons described in the previous section.

### C12ORF57/Grcc10 loss of function causes a decrease in the expression of pCREB, cFOS and ARC

Because the loss of C12ORF57 disrupts synaptic AMPA downscaling and current, we investigated whether there were changes in the negative regulation of AMPA synapses involving CREB and ARC signaling. This is because phosphorylated CREB (p-CREB) regulation of the immediate early gene, *Arc*, is implicated in the synaptic downscaling of AMPA receptors and the regulation of surface GluA2 expression in response to neuronal excitation. ^1,17,26–30^ To determine whether this pathway is altered in our KO mice, we stained primary hippocampal *Grcc10* WT and KO neurons isolated at E18 at 20 days *in vitro* using an antibody that binds specifically to the Ser133 p-CREB and ARC and measured fluorescence intensity. We observed that KO neurons express a significantly lower level of Ser133-pCREB (-29.80±8.55%, (p=0.0008, t=3.66, df=38, N=20, Welch’s T-Test) and ARC (-65.79±9.18%, p<0.0001, t=7.17, df=38 N=24, Welch’s T-Test) compared to WT neurons (**Figure 4A-C**). These data were also supported by western blot findings of decreased pCREB in whole brain lysates of KO mice compared to WT (-31.26±12.47%, (p=0.024, t=2.5, df=14.72, N=5, Welch’s T-Test) and decreased intermediate early genes ARC (- 48.03±8.44%, p=0.0036, t=5.69, df=4.36, N=5 Welch’s T-Test) and cFOS (-49.94%±10.45, =0.0043, t=4.78, df-5.288, N=5, Welch’s T-Test) (**Figure 4D-F**). These results suggest that *Grcc10* loss of function leads to a decrease in CREB phosphorylation and decreased expression of ARC and cFOS.

**Figure 4.**
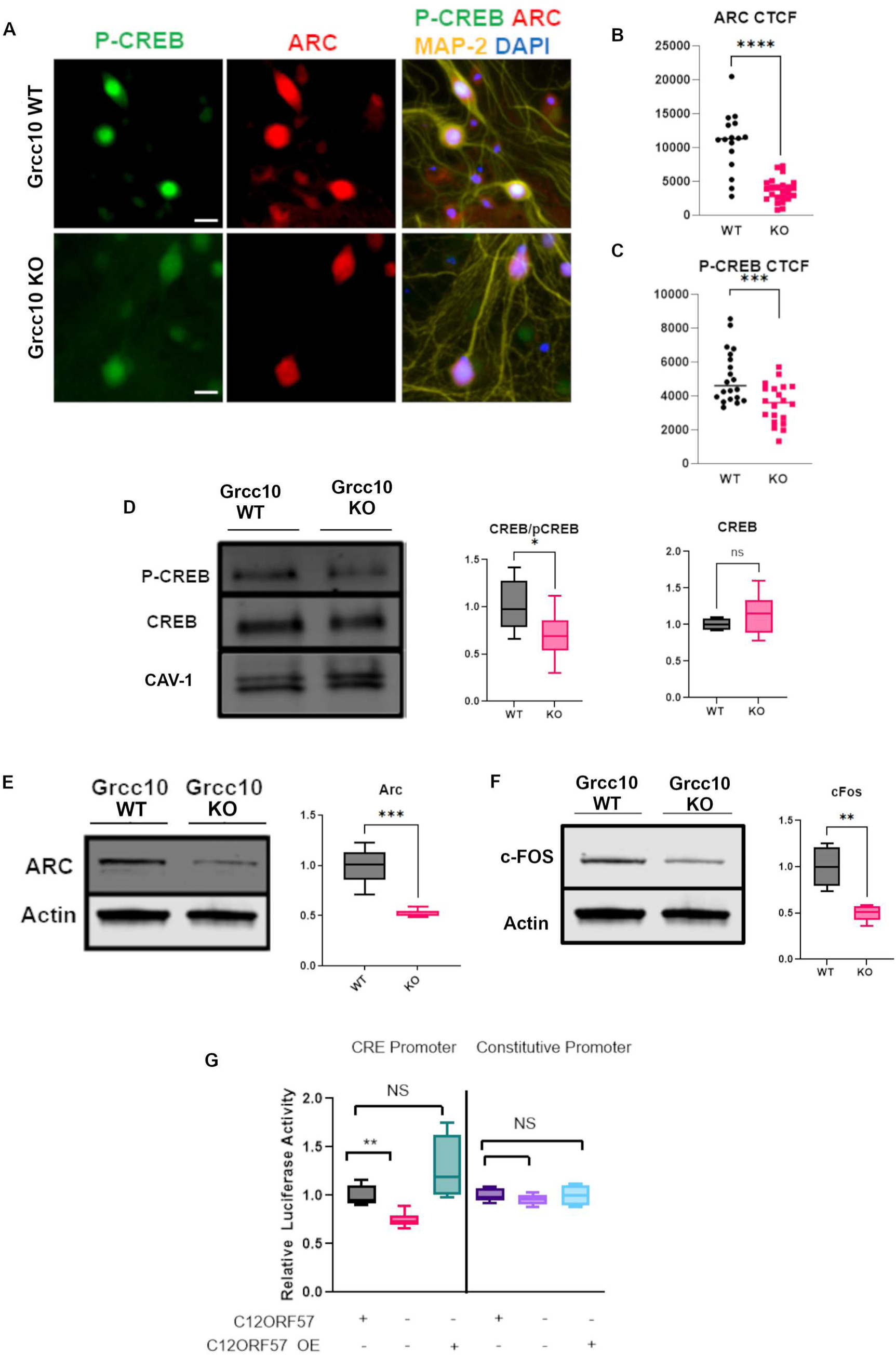
Loss of C12ORF57 decreases levels of downstream CAMK4 targets. **A)** IHC staining of primary hippocampal neurons from *Grcc10* (upper panels) and *Grcc10* KO (lower panels) mice for phospho-CREB (green, left), ARC (red, center) with overlay of MAP-2 Yellow and DAPI (blue, right). Scale bar 5µm **B and C)** Fluorescent quantification of ARC (**B**) (p=0.0008, N=20 for each genotype, Welch’s T-Test) and p-CREB **(C)** (p<0.0001, N=24, Welch’s T-Test). Median RFU are indicated by horizontal line. **D) Left:** Western blot of whole brain lysates of pCREB (top) CREB (middle) and Caveolin-1 (Cav-1) loading controls (bottom). **Middle**: Ratio of CREB/pCREB fluorescence relative to Cav-1 loading control (p=0.024, N=10, Welch’s T-Test). Normalized to WT Mean. **Right:** Total CREB normalized to loading control (p=0.29,N=5, Welch’s T-test) **E) Left:** Western blot of whole brain lysates from *Grcc10* WT and *Grcc10* KO mice with anti-ARC (top) and anti-actin control (bottom) antibodies. **Right:** ARC band intensity to actin normalized to mean WT intensity (p=0.0036, N=5 for each genotype, Welch’s T-Test). **F) Left:** Western blot of whole brain lysates from *Grcc10* KO and WT mice with anti-cFos (top) and anti-actin control (bottom) antibodies. **Right:** Ratio of cFos band intensity to actin normalized to WT mean (p=0.0043, N=5 for each genotype) **G)** Relative luminescent signal from Renilla CRE luciferase assay on HEK293 cell lysates normalized to WT control (N=6 for each genotype, Welch’s T-Test.) and constitutive promoter positive controls (right). For all box and whisker graphs, min max and median and interquartile range are represented. Statistical significances are indicated by the following symbols \**P* ≤0.05, \*\**P* ≤0 .01, \*\*\**P* ≤ 0.001, \*\*\*\**P* < 0.0001.

To further study the effect of *C12ORF57*/*Grcc10* loss of function on CRE-dependent transcriptional activation, we performed a dual luciferase assay as described by Bellis et al. ^31^ We transfected both *C12ORF57* WT and KO HEK293T cells with one construct containing *Renilla* luciferase as a reporter of CRE promoter activity and another one expressing firefly luciferase under a constitutive promoter as a control. We observed that KO cells had a significant decrease in normalized CRE-driven *Renilla* luminescence compared to WT cells (-25.40±3.78%, ANOVA, F=20.74, p<0.0001, Tukey post-hoc test, p=0.019, N=3 per experimental condition). Moreover, transfection of *C12ORF57*-/- cells with C12ORF57 restored CRE-driven luciferase signal to wildtype levels (+27.56±17.10%, p=0.12, Tukey post-hoc test) (**Figure 4G**). These data show that loss of C12ORF57 reduces CRE-dependent transcriptional activation, potentially impacting the negative regulatory elements upstream of AMPA, CREB and ARC.

### C12ORF57 enhances CAMK4 activity through interactions with its autoinhibitory domain

The lack of significant homology of GRCC10 to any known protein impaired our ability to predict its cellular mechanisms *a priori*. To bypass this limitation, we analyzed the GRCC10 protein interactome using two independent unbiased Yeast 2-hybrid screens utilizing mouse brain libraries.^32^ Both unbiased genome-wide screens resulted in a single positive clone that encoded the calcium calmodulin associated kinase IV (CAMK4). We conducted direct co-immunoprecipitation of GRCC10 from whole WT and KO mouse brain lysate and confirmed its interaction with CAMK4 in WT lysate which was abolished in KO lysate (**Figure 5A**). Discovery of C12ORF57’s interaction with CAMK4 is particularly informative because CAMK4 is a known upstream regulator of both CREB and ARC and plays a role in synaptic downscaling. ^1,28,33^

**Figure 5.**
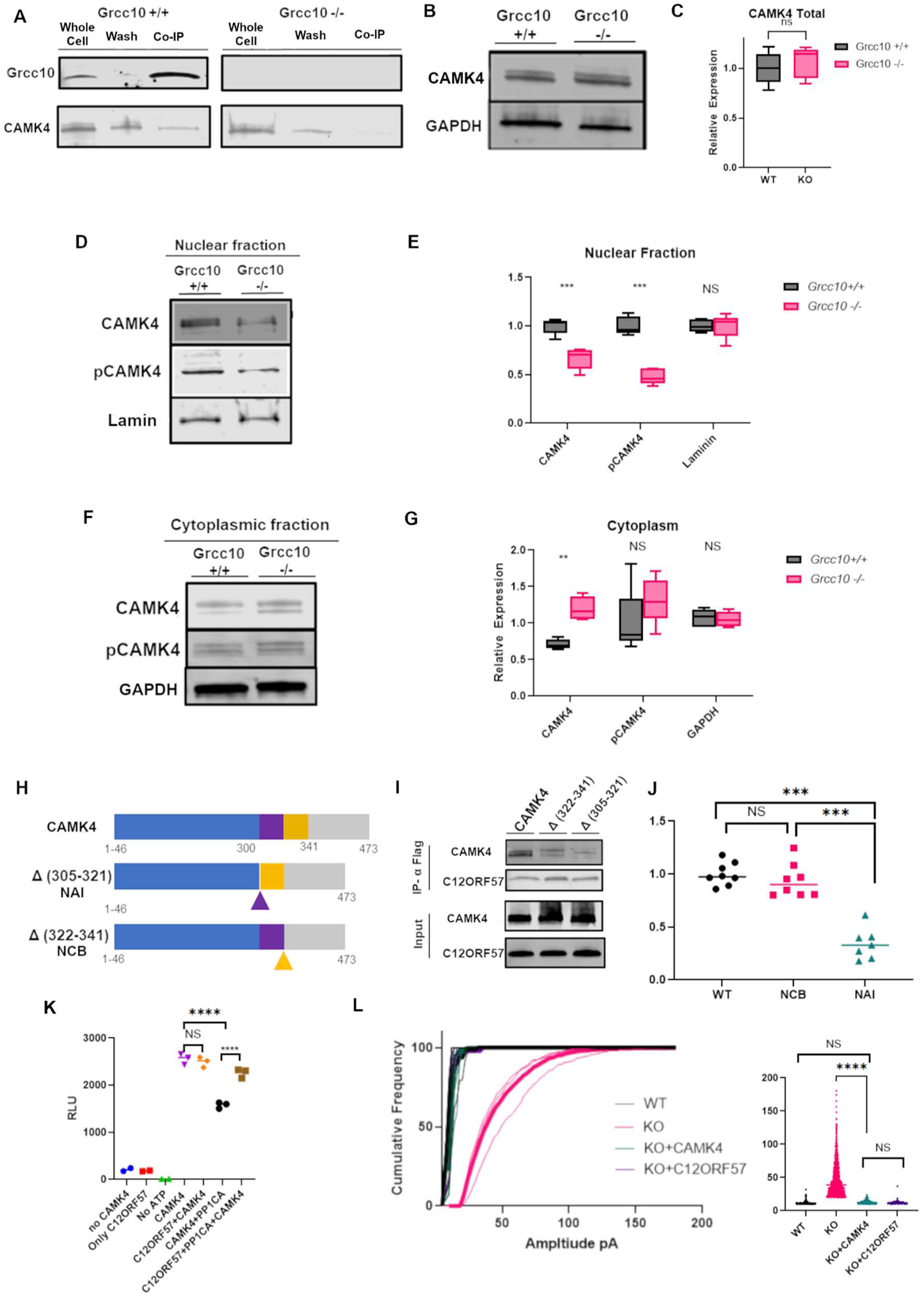
C12ORF57 binds CAMK4 and modulates its phosphorylation and activity. **A)** Western blot showing co-immunoprecipitation of CAMK4 with C12ORF57 from *Grcc10+/+* whole brain lysate (left) with *Grcc10-/-* brain negative control (right) showing no GRCC10 expression or CAMK4 pulldown **B)** Representative CAMK4 and GAPDH western blot bands from brain whole cell lysates of *Grcc10 +/+* and *Grcc10-/-* mice **C)** Mean relative CAMK4 western blot band intensity normalized to WT (p=0.34, N=6 for each genotype Welch’s T-Test) **D)** Representative CAMK4, pCAMK4 and Lamin A/C (control) bands on western blot from nuclear fraction of *Grcc10* WT and KO whole brain lysates **E)** Relative CAMK4, pCAMK4 and Lamin A/C band intensity normalized to *Grcc10 WT* levels. (p=0.001, N=6 for each genotype, 2-way ANOVA with Sidak’s multiple comparison test). **F)** Representative CAMK4, pCAMK4 and GAPDH (control) bands on western blot from cytoplasmic fractions of whole brain **G)** Relative CAMK4 pCAMK4 and Lamin band intensity normalized to *Grcc10* WT levels (p=0.016, N=6 for each genotype, 2-way ANOVA, Sidak’s multiple comparison test) in KO compared to WT. **H)** Schematic of WT CAMK4 (top) with kinase domain (blue), calmodulin binding domain (purple) and autoinhibitory/regulatory domain (yellow). Construct Δ(322-341) which lacks the autoinhibitory domain (NAI, middle) and construct Δ(305-321) which lacks the calmodulin binding domain (NCB, bottom) **I)** Representative western blot of co-IP of CAMK4 constructs with C12ORF57-FLAG **J**) Relative CAMK4 band intensity between WT, NCB (p=0.24, N=8 replicates for each construct, ANOVA) and NAI (p<0.0001, N=8 replicates for each construct, ANOVA) CAMK4 constructs. **K**) Relative luminescence from *in vitro* CAMK4 kinase activity assay for C12ORF57, CAMK4, CAMK4+C12ORF57, CAMK4+PP1CA, CAMK4+PP1CA+C12ORF57, no CAMK4, and no ATP (N=3 per condition, ANOVA). **L**) **Left:** Graph of cumulative frequency (x-axis) of mEPSC against amplitude (pA) in DIV 17 pyramidal neurons including CAMK4 transfected control (green) KO (pink) WT (black), and C12ORF57 transfected (purple). The mean curve of each group is bolded. **Right:** Amplitude of WT, KO, CAMK4 transfected (KO+CAMK4) and C12OR57 transfected (KO+C12ORF57) AMPA mEPSCs (p<0.0001, N=6 for each group, ANOVA). For all box and whisker graphs, min, max and median and interquartile range are represented. Statistical significances are indicated by the following symbols \**P* ≤0 .05, \*\**P* ≤0 .01, \*\*\**P* ≤ 0.001, \*\*\*\**P* < 0.0001.

Given the close association of C12ORF57/GRCC10 with CAMK4, we next investigated whether it has a role in modulating CAMK4 activity. In prior studies, phosphorylated CAMK4 has been shown to be largely confined to the nucleus.^34,35^ This localization for CAMK4 was shown to be mediated by phosphorylation at amino acid (AA) Thr200 in humans and via Thr196 in mice. ^30,36,37^ We measured the expression of CAMK4 in *Grcc10* WT versus KO brains by western blot and observed that differences between total CAMK4 were not significant between WT and KO brains (-0.74 ±0.73%, p=0.34, t=1.01, df=6.97, N=6 Welch’s T-Test). However, we did see a shift in CAMK4 distribution within the nucleus of KO brains compared to WT brains with significantly less pCAMK4 (-66.66±6.83%, p=0.0037, F=18.33, df=3, N=5, ANOVA with Tukey) and total CAMK4 (-48.1±6.73%, p=0.002 F=14.89, N=5, df=3, ANOVA with Tukey post-hoc test). We saw the opposite shift in cytoplasmic fraction, where KO brains showed an increase in CAMK4 (+31.6±20.69%, p=0.0035 2-way ANOVA, F=5.50, DF=2, N=6) compared to WT, and a similar, though not statistically significant, decrease in pCAMK4 (+31.87±19.51, p=0.1848, N=6, Tukey post-hoc test) (**Figure 5B-G**). These results suggest that loss of C12ORF57 disrupts the balance of nuclear versus cytoplasmic CAMK4, with increased nuclear phosphorylated CAMK4 in WT brains when compared to KO brains.

To further probe how C12ORF57/GRCC10 interacts with CAMK4 and because we observed changes in CAMK4’s phosphorylation and localization with C12ORF57 loss, we investigated the roles of various regulatory domains of CAMK4. We created two modified forms of CAMK4 constructs lacking distinct regulatory domains: one which lacks the autoinhibitory domain (AA 305-321) (NAI), which binds CAMK4-negative regulatory proteins phosphatase 1 and 2 (PP1 and PP2), and the other lacking the calmodulin binding domain (AA 322-341) (NCB) (**Figure 5H**). We co-transfected these constructs in HEK293T cells with a flag-tagged version of C12ORF57 and performed a co-immunoprecipitation assay with CAMK4. We observed that the deletion of the autoinhibitory domain significantly reduced the ability of CAMK4 to bind to C12ORF57-FLAG (-65.76±6.76%, p<0.0001, N=8, ANOVA with Tukey post-hoc test) when compared to wild type CAMK4, whereas deletion of the calmodulin binding domain resulted in no significant change in binding (-6.13±6.81%, p=0.66, ANOVA with Tukey post-hoc test, N=8) (**Figure 5H-J**). This suggests that the CAMK4 autoinhibitory domain plays a significant role in its interaction with C12ORF57.

The association of C12ORF57 with CAMK4’s autoinhibitory domain that interacts with protein phosphatases, along with the decrease in pCAMK4 in KO brain nuclei, led us to hypothesize that C12ORF57/GRCC10 disrupts the activity of phosphatases that modulate CAMK4 activity. To investigate this possibility, we measured *in vitro* CAMK4 kinase activity in the presence or absence of C12ORF57 and/or serine/threonine-protein phosphatase alpha catalytic subunit 1 (PP1CA) using an *in vitro* kinase enzyme system that enables the quantification of ATP consumption by CAMK4 via luminescence. ^33^ We found that the addition of PP1CA significantly decreases CAMK4 *in vitro* activity by 39.32% (-981.00±75.95 Relative luminesce units, RLU, p=<0.0001, ANOVA with Tukey’s post-hoc test), which was expected due its negative regulatory activity *in vivo* (**Figure 5K**). We also found that adding C12ORF57 does not affect CAMK4 kinase activity compared to CAMK4 alone (-66.33±90.83 RLU, p=85, F=448.3, df=6, N=3,p=0.95, ANOVA with Tukey’s post-hoc test). The presence of both C12ORF57 and PP1CA caused CAMK4 activity to decrease significantly than just PP1CA alone (-681.00±67.39 RLU p<0.0001, ANOVA with Tukey’s post-hoc test) (**Figure 5K**). These findings, in conjunction with our previous data, suggest that C12ORF57 enhances CAMK4 activity by diminishing the efficacy of phosphatase inhibition by binding to the autoinhibitory domain of CAMK4 and increasing the relative amount of active phosphorylated CAMK4.

*Overexpression of CAMK4* is sufficient to rescue *Grcc10* excitability phenotype.

Since we hypothesize that CAMK4 is downstream of C12ORF57/GRCC10, we tested whether overexpression of CAMK4 would be sufficient to rescue the increased AMPA mEPSC phenotype in *Grcc10-/-* neurons. We performed whole cell recordings on isolated excitatory hippocampal neurons from embryonic day 18, *Grcc10* WT, and KO mice on DIV 14-18 that were transfected with a lentivirus carrying a *Camk4-Pp2A-Gfp* fusion under *CMV* promoter control versus a *GFP* vector as a negative control and a *C12ORF57* positive control. *Grcc10* KO neurons transfected with *CAMK4* had AMPA amplitudes statistically significantly lower (p<0.0001, F=240.8, dF, 3,N=6, ANOVA) than vector-infected KO neurons (12.54±2.29 pA vs. 46.11± 20.70 pA, respectively, p<0.0001, Tukey post-hoc test) and equal to WT neurons (12.30±3.52 pA p=0.99, N=6, Tukey post-hoc test) and *C12ORF57*-transfected KO neurons (12.55±3.684pA, p=0.9987, N=6, Tukey post-hoc test) (**Figure 5L**). These data suggest that CAMK4 is indeed downstream of C12ORF57 and that the overexpression of CAMK4 is sufficient to overcome the loss of C12ORF57 function and rescue the AMPA amplitude phenotype.

## Discussion

In this paper, we report the generation and characterization of a GRCC10 knockout mouse model of Temtamy syndrome. Our *Grcc10* knockout mice display similar brain phenotypes (seizures, shortened corpus callosum) to those observed in human Temtamy syndrome, caused by mutations in the closely related and poorly characterized human ortholog, *C12ORF57*. Due to the lack of recognized protein motifs or domains in C12ORF57, our *Grcc10* knockout mice provide a novel window to elucidate the precise molecular function of these undescribed genes.

Based on our observations, we propose a model where C12ORF57 protects CAMK4 from dephosphorylation and bolsters its phosphorylation life-time, thereby enabling C12ORF57’s homeostatic role in reining in excitatory synaptic up-scaling in excitatory neurons (**Figure 6**). In this model, the loss of C12ORF57 leads to a decrease in the proportion of active, phosphorylated, CAMK4, reducing negative feedback of excitatory synaptic scaling and increased AMPA expression at synapses. These events lead to higher baseline neuronal excitability and increased occurrence of seizures seen in both TS patients and our murine knockout model. Because sequence analysis of human and murine CAMK4 shows identical calmodulin and autoinhibitory domains, as well as 98.3% amino acid identity between these orthologs, it is likely that our biochemical findings in mice would generalize to humans. It should be noted that this is not the first time CAMK4 activity has been implicated in epilepsy in humans; disruptions involving a regulatory miRNA and CAMK4 point mutations have been reported in previous clinical cases. ^38–40^ The regulation of the phosphorylation state of CAMK4 would also represent a novel mechanism by which individual neurons could tune their baseline excitability dynamically and may underlie how individual neurons can have variable responses to the same electrical stimulus.

**Figure 6.**
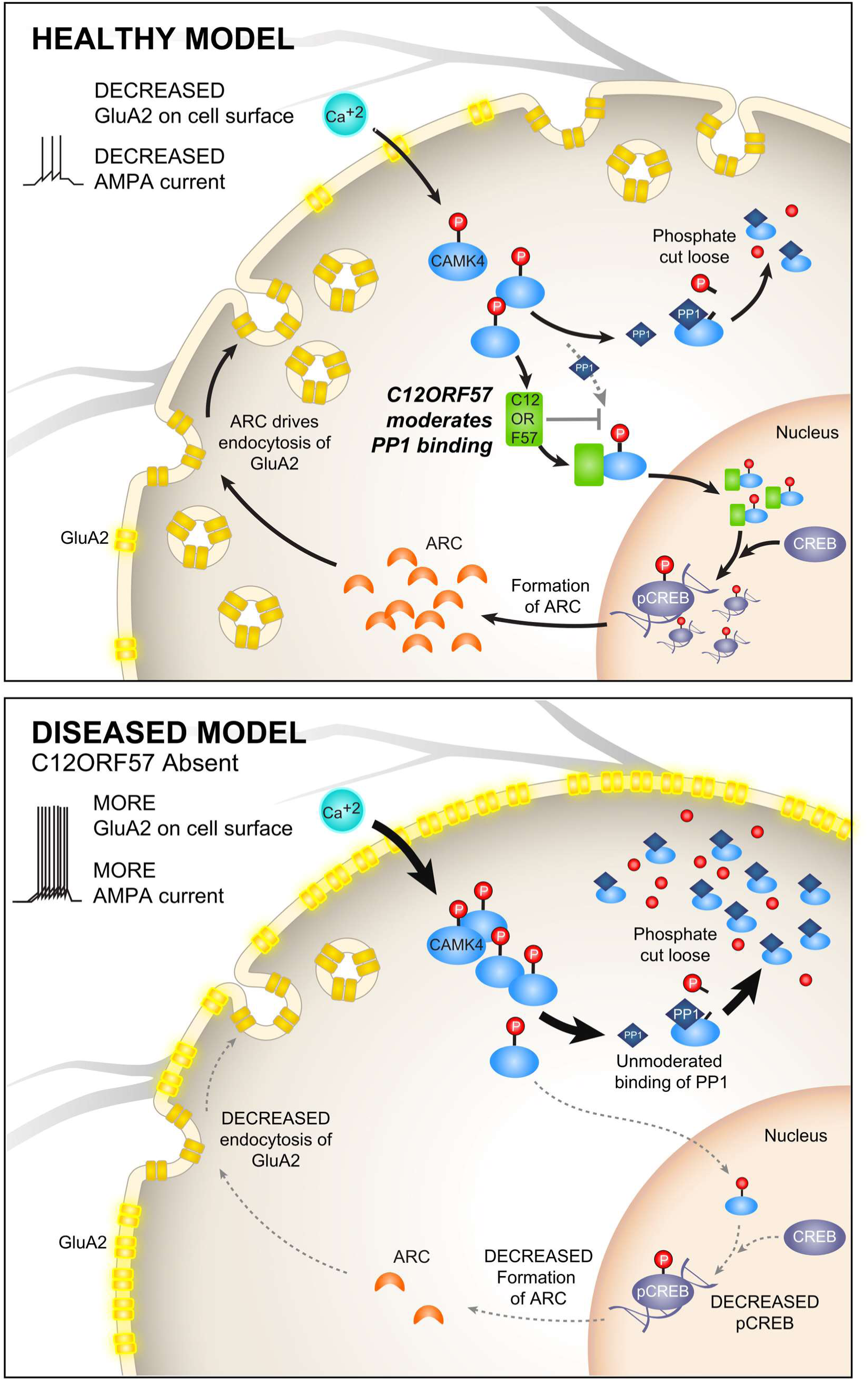
Proposed Model of C12ORF57 Function. The proposed normal function of C12ORF57 (top panel). Under normal conditions C12ORF57 binds CAMK4 and blocks dephosphorylation leading to increased active CAMK4 and activation of ARC and down regulation of AMPA. Loss of C12ORF57 (bottom), where in response to the same level of calcium ion influx during depolarization/presynaptic excitation, more CAMK4 is dephosphorylated, and less is able to translocate to the nucleus, decreasing downstream pCREB transcription and ARC transcription resulting in unopposed long-term potentiation, increased GluA2 on cell surface and increased AMPA current.

We demonstrate that loss of *Grcc10* occluded the normal AMPA GluA2 downscaling in response to GABAR inhibition. We also showed that loss of *Grcc10* decreased known excitatory synaptic regulators CREB, Arc and cFos, and that at baseline *Grcc10 -/-* neurons express greater levels of AMPA and have increased mEPSCs. This suggests that C12ORF57 is capable both of regulating the baseline excitatory potential of neurons as well as the dynamic response of neurons to overexcitation, acting as a neuronal excitatory “rheostat” whose loss leads to an unexpected and dramatic increase in AMPA current and signaling. While the full mechanism and scope of this rheostat function will require further study to elucidate, we believe that CAMK4, the binding partner we discovered, provides a plausible mechanism.

The fine tuning of AMPA current would require a protein regulator, which is itself tightly and dynamically regulated. The activation of CAMK4 is a transient and tightly regulated event that requires both Ca^2+^/calmodulin binding and phosphorylation of the kinase at residue Thr200 by an upstream CaMK kinase (CaMKK). Upon phosphorylation CAMK4 trafficsinto the nucleus. ^33^ Expression of either dominant negative or constitutively active forms of CAMK4, respectively scales up or down excitatory synaptic strength, and disrupts the ability of cells to dynamically respond to stimuli. ^16,33,41^ As anticipated, a disruption in CAMK4 activity led to a destabilization of neuronal homeostasis in our *in vitro* murine hippocampal excitatory neuronal model. Loss of *Grcc10* increased AMPA mEPSCs and GluA2 expression and the inability to downregulate excitatory GluA2 in response to increased activity upon GABAR blockade, as observed with CAMK4 dominant negative mutations. Furthermore, we observed a decrease in the ratio of nuclear pCAMK4/CAMK4 in our mouse model similar to changes in this ratio reported with the overexpression of a dominant negative version of CAMK4.^44^ Our *in vitro* data also supports the observation of decreased pCAMK4 with the loss of C12ORF57 decreasing downstream pCAMK4 signaling through pCREB and ARC. CREB acts upstream of the immediate-early gene *ARC* which in turn is responsible for downscaling excitatory synapses in response to neuronal excitation. ^30,42^ CREB and ARC signaling has been shown to increase endocytosis of surface GluA2 expression in response to increased calcium flux from neuronal excitation ^1,17^. Prior published evidence showed that ARC is involved in synaptic scaling by interacting with endophilin to enhance endocytosis of AMPA receptors, which is crucial for neuronal excitatory/inhibitory homeostasis.^17,26–29,43^ CAMK4 is known to regulate the activity of CREB by phosphorylating its Ser133 residue.^34,44,45^

Previous studies have shown that the autoinhibitory domain of CAMK4 stably associates with protein serine/threonine phosphatases 1 and 2A (PP1; PP2A) to negatively regulate its kinase activity.^33,46^ These interactions modulate CAMK4’s role in homeostatic excitatory synaptic scaling, a mechanism by which neurons adapt their synaptic strength through upregulation or downregulation of surface expression of AMPA receptors. ^16,27,46^ We noted that C12ORF57 binds CAMK4 and the presence of the autoinhibitory phosphatase binding domain was important in maintaining this interaction whereby C12ORF57 antagonized the PP1CA-mediated decrease in CAMK4 activity *in vitro*. These results give a further plausible mechanism by which C12ORF57 could regulate excitatory synaptic strength through modulation of CAMK4 activity by slowing its dephosphorylation and thereby prolonging pCAMK4 signaling.

A key question arising from our model is why previous studies using CAMK4 knockout mice did not observe a similar seizure phenotype.^47^ It is possible that loss of C12ORF57 in excitatory neurons and a subsequent decrease in the pCAMK4/CAMK4 ratio creates a functional CAMK4 knockdown and that cells do not tolerate knockdown as well as a knockout or complete loss of CAMK4 function; that CAMK4 is sensitive to changes in both subcellular localization as well as relative levels when compared to other CaM kinases bolsters this hypothesis.^34,48,49^ In neuronal subsets such as excitatory neurons that have *high* levels of endogenous CAMK4, the levels of other CaMKs such as CAMKI are the principal regulators of neuronal firing, while in neurons with *low* endogenous CAMK4 levels, CAMK4 is the primary regulator of neuronal firing^50^ A complete knockout would react distinctly from a knockdown as it would would cause the cell to rely on other compensatory pathways causing CAMK4 to be no longer limiting. For these reasons we propose that our data suggests that it is the relative levels of phosphorylated CAMK4 that cause disruptions in neuronal excitatory homeostasis which are not captured in previous complete knockout experiments.

We acknowledge the simplified nature of our model, and that further investigation is needed to fully elucidate the precise impact of C12ORF57 on the kinetics, localization, and phosphorylation of various CaM kinases. Also, while we saw no increased NMDA or GABA activity in *Grcc10* KO neurons, we cannot fully determine whether this is due to changes in presynaptic release probability versus purely increased postsynaptic excitation/detection, specifically its effect on AMPA current frequency. Further work is necessary to determine what, if any, effect C12ORF57 has on presynaptic transmission. Additionally, the high perinatal mortality makes it difficult to study certain aspects of the knockout phenotype and limits broader behavioral characterization. The development of a conditional knockout would be helpful to further studying the phenotypic features found in human patients, such as optic coloboma, motor delay, social impairment, and epilepsy, since complete null mice cannot tolerate EEG placement and monitoring. Another area for future study is that while most mutations currently discovered in human patients involve a loss of function of the start codon, there are several missense, nonsense and splice site mutations reported in compound heterozygotes such as p.L51Q mutations.^6,11^ Patients with non-initiator methionine mutations have similar phenotypes and severity (epilepsy, agenesis of the corpus callosum and optic coloboma), suggesting that these mutations also result in loss of function of the protein.^4,6,8^ Further research into specific missense mutations may provide additional insight into key amino acid residues and domains within the C12ORF57 protein.

In conclusion, we have found that the novel *Grcc10* knockout mouse shares many phenotypic features with Temtamy syndrome patients, making it a suitable model in which to study the function of its human ortholog, *C12ORF57*. We have shown through multiple lines of evidence in murine, human cellular, and *in vitro* models that – a) C12ORF57 binds CAMK4, b) C12ORF57 loss decreases CAMK4 phosphorylation and increases mEPSC activity, and c) C12ORF57 loss decreases downstream signaling through the synaptic scaling pathway of CREB, ARC, and GluA2. These findings provide the first evidence for the molecular function of C12ORF57 in neuronal development and homeostasis, as well as a possible explanation for the pathogenesis behind some of the phenotypic features of Temtamy syndrome. Further studies, using conditional knockout models, would shed further light on additional functions of this protein and elucidate its precise role in synaptic homeostasis. These studies have the potential to provide critical insight into how C12ORF57 regulates neuronal excitatory synaptic scaling and protects neurons from excessive excitability and maintains neuronal homeostasis.

## Acknowledgments

We thank the staff of the University of Queensland Biological Resources (UQBR) animal facility and the QBI Advanced Microscopy and Histology Facilities, for their expertise and assistance in this project.

Microscopy support was provided by the Center for Advanced Microscopy and the Nikon Imaging Center at UCSF

Suling Wang for providing the illustrations in Figure 6 and Figure 1 Panel 1D.

We would like to thank Dr. Saleem Alkeraye of the College of Medicine, King Saud University, Riyadh Saudi Arabia for their assistance in obtaining C12ORF57 patient skin biopsy.

We would like to thank the multiple families and patients who provided samples and participated in our research

## Funding Sources

Research in this project was funded by NINDS RO1 NS0587221-15, NEI EY02162-39, EY032197 and NHMRC Principal Research Fellowship (GNT1120615).

## Competing Interests

The authors declare no competing interests.

## Notes

### Competing Interest Statement

The authors have declared no competing interest.

### Summary of Updates

Addition of new author Kok Siong Chen and updates to abstract, discussion and introduction

