## Supplemental Figures for "C12ORF57: a novel principal regulator of synaptic AMPA currents and excitatory neuronal homeostasis"

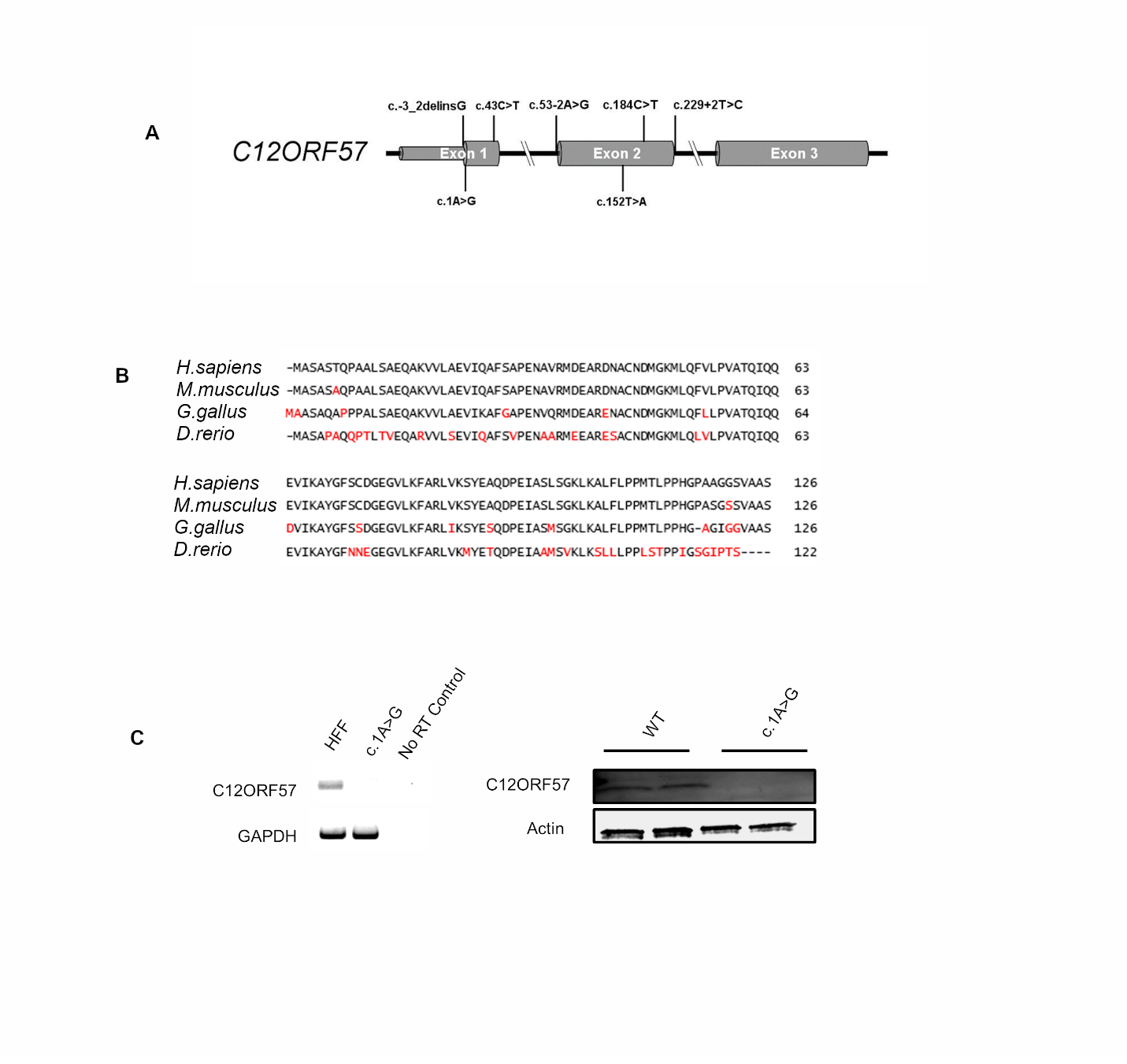

**Supplemental Figure 1** *C12ORF57 Well conserved throughout evolution* **A)** Gene schematic of all published mutations in C12ORF57 **B)** Protein sequence alignment between Homo *sapiens,* Mus *musculus,* Gallus *gallus,* and Danio *rerio.* Amino acid differences are marked in red and sequences without homology are marked with dashes. **C)** RT PCR (Left) and western blot on whole cell lysate (Right) from primary fibroblasts from patients with C12ORF57 1.A>G mutations.

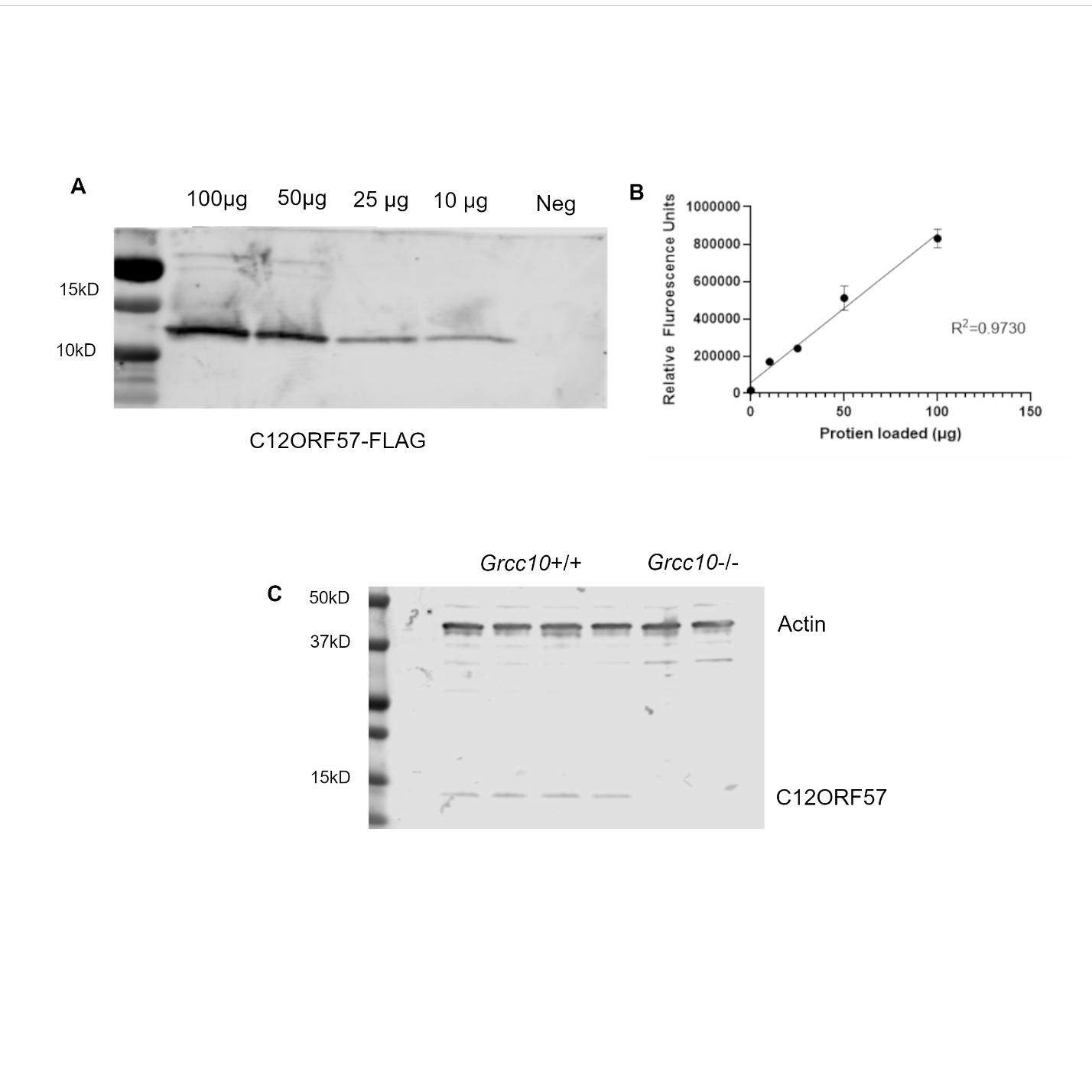

**Supplemental Figure 2** *C12ORF57 Rabbit Antibody is specific in western blots* A) western blotting on HEK293 lysates (total protein loaded per lane labeled above) transfected with C12ORF57-FLAGx2 tagged protein shows bands at expected molecular weight (17 KD) and no band in untransfected control. **B)** Graph showing relative fluorescence units against total protein loaded per lane, showing robust linear (R^2^=0.9730) correlation **C)** Whole brain lysates of Grcc10 +/+ mice show staining on Western blot with Anti-C12ORF57 antibody expected C12ORF57 bands expected size at 13 kD and no bands in Grcc10 -/- mice.

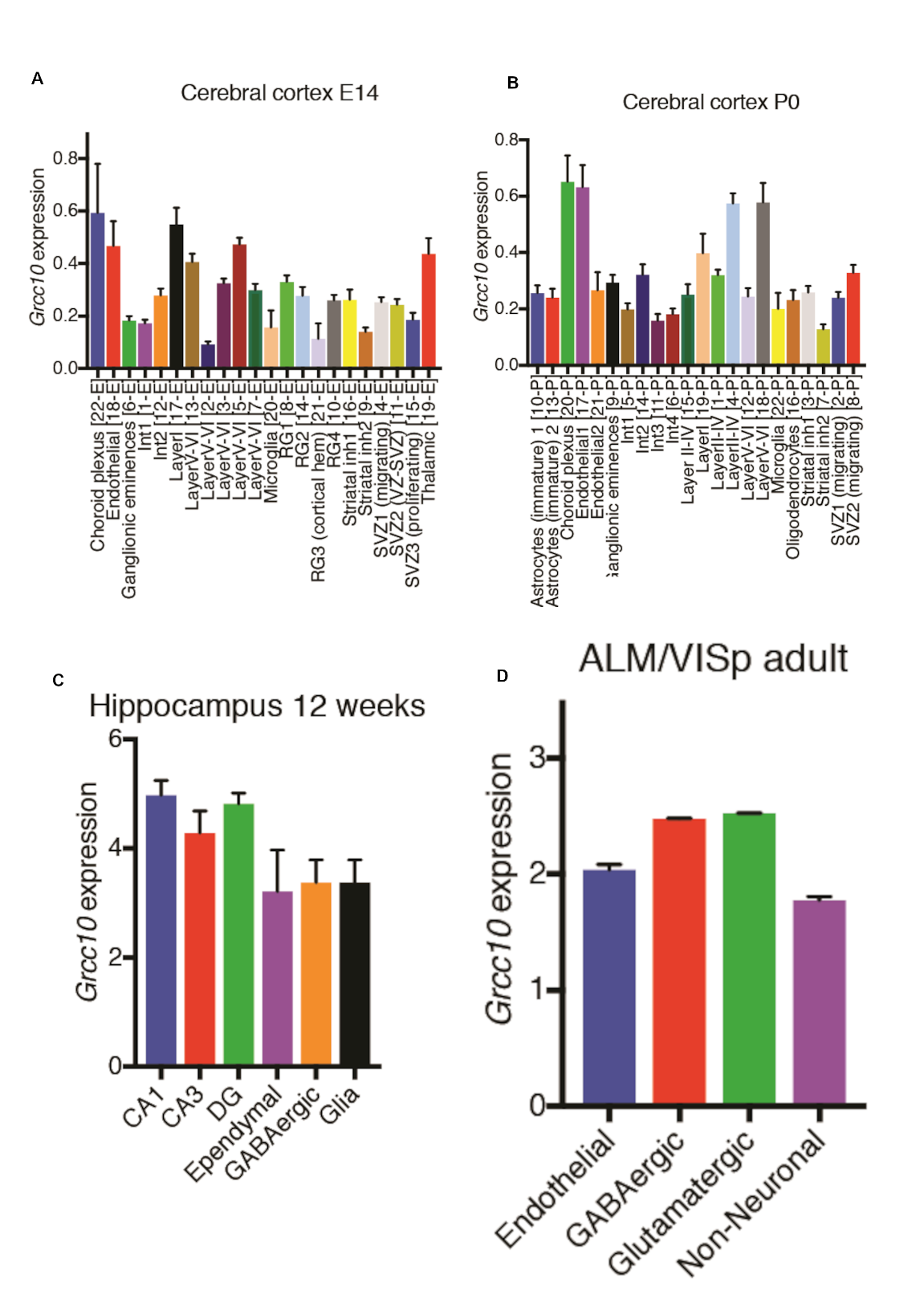
**Supplemental Figure 3** *C12ORF57/Grcc10 Single Cell Expression* Grcc10 expression mapped across cerebral cortex single cell data at **A)** E14 and **B)** P0 from Loo et al 2019 (52). **C)** Cell type specific expression in mouse hippocampus at 12 weeks of age from Habib et al 2017 (53) **D)** Expression in cell type specific from adult V1 and anterior lateral motor cortex from Allen Brain Atlas data sets. All error bars represent SD.

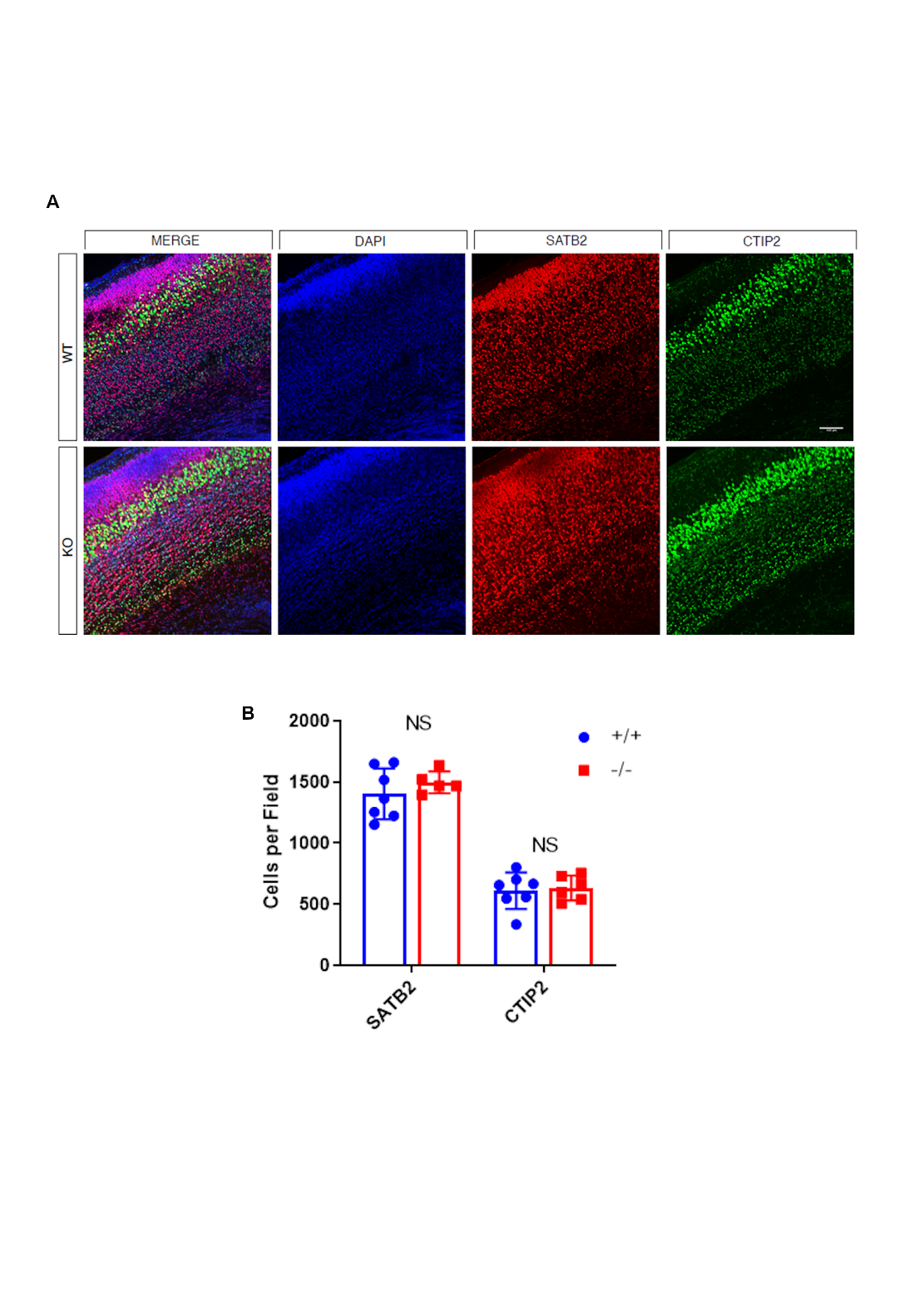

**Supplemental Figure 4** *Loss of C12ORF57 does not disrupt overall cortical layering*

**A)** Post natal day 0 mouse cortical slices were of *Grcc10* WT (top) and KO (bottom) were stained for Layer II/II marker SATB2 (Red) (p=0.36, Welch’s T test) and Layer IV/V marker CTIP2 (Green) (p=0.76, Welch’s T-Test) with DAPI (blue) co-staining. **B)** Mean number of cells SATB2 cells (Right) and CTIP 2 Cells (Left) per HPF. All error bars represent SEM.

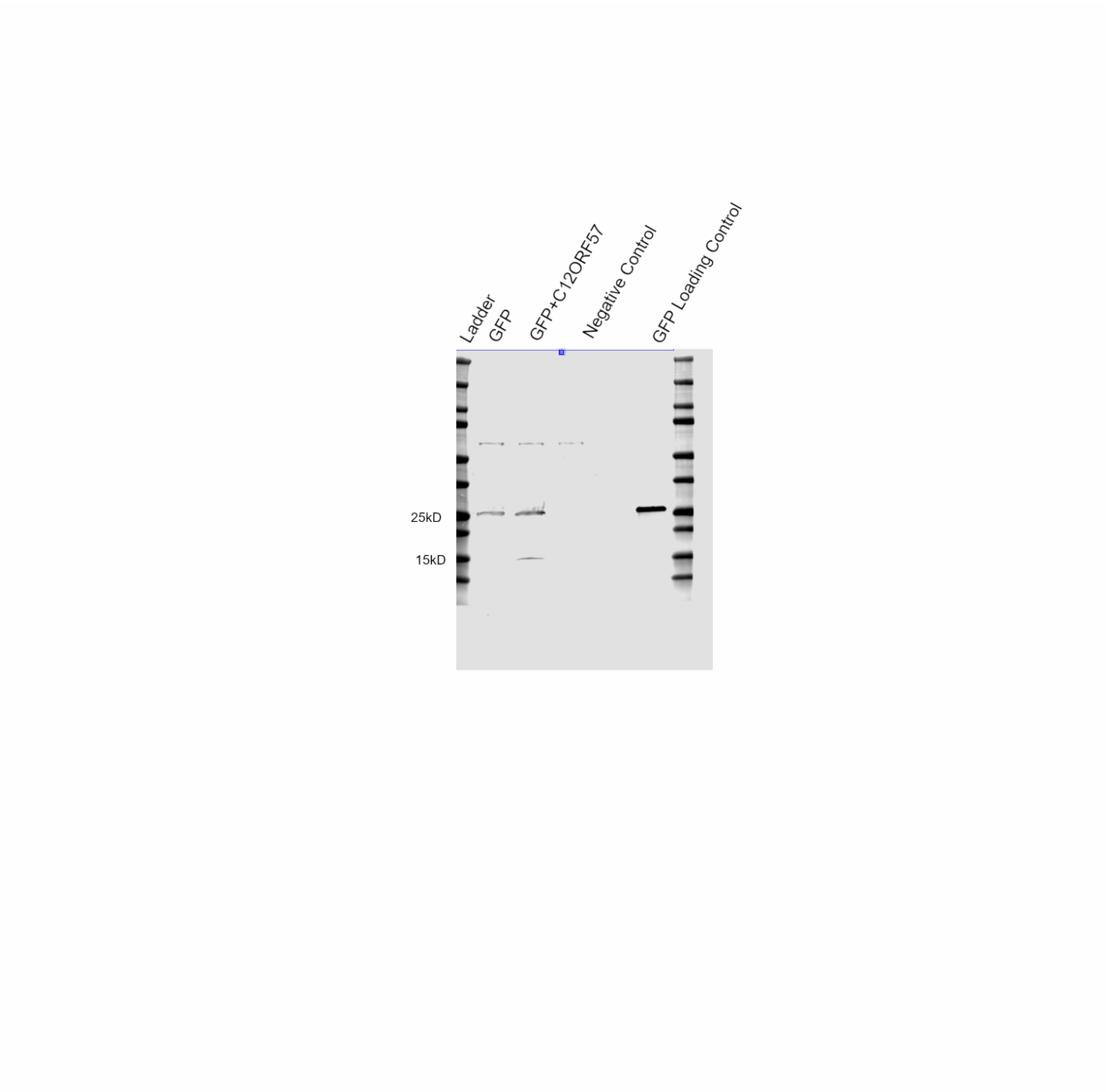

**Supplemental Figure 5** *Lentiviral* *C12ORF57-P2A-GFP* Western blot of whole cell lysate of HEK293T cells transfected with GFP under Syn promoter control (leftmost lane), vector C12ORF57-P2A self-cleaving peptide-GFP construct (second lane), untransfected HEK293T cells (third lane), and 1 ug of purified GFP positive control (right lane).

**Supplemental Video 1** *Observed Clonic Tonic Seizure in Grcc10-/- Mouse Pup*

Video of spontaneous tonic-clonic movements suggestive of seizure observed during a routine care check in a P18 *Grcc10-/-* mouse.
